# SIMON, an automated machine learning system reveals immune signatures of influenza vaccine responses

**DOI:** 10.1101/545186

**Authors:** Adriana Tomic, Ivan Tomic, Yael Rosenberg-Hasson, Cornelia L. Dekker, Holden T. Maecker, Mark M. Davis

## Abstract

Machine learning holds considerable promise for understanding complex biological processes such as vaccine responses. Capturing interindividual variability is essential to increase the statistical power necessary for building more accurate predictive models. However, available approaches have difficulty coping with incomplete datasets which is often the case when combining studies. Additionally, there are hundreds of algorithms available and no simple way to find the optimal one. Here, we developed Sequential Iterative Modelling “OverNight” or SIMON, an automated machine learning system that compares results from 128 different algorithms and is particularly suitable for datasets containing many missing values. We applied SIMON to data from five clinical studies of seasonal influenza vaccination. The results reveal previously unrecognized CD4^+^ and CD8^+^ T cell subsets strongly associated with a robust antibody response to influenza antigens. These results demonstrate that SIMON can greatly speed up the choice of analysis modalities. Hence, it is a highly useful approach for data-driven hypothesis generation from disparate clinical datasets. Our strategy could be used to gain biological insight from ever-expanding heterogeneous datasets that are publicly available.

## Introduction

The immune system is comprised of multiple cell types that work together to develop an effective response to a given pathogen. However, which of these myriad cell types are important in a particular response is not well understood. The increasingly common systems immunology approach measures gene expression and different cells and molecules in the immune system during an infection or vaccination and uses computational methods to discern which components are most important (1–6). These studies have the practical goal of determining what makes one vaccine formulation better than another or how individuals vary. In addition, it may suggest a mechanistic understanding of how an effective immune response is achieved. To accomplish this, an accurate modeling of the complex processes that lead to a successful outcome is crucial.

Over the past few years, many systems studies of influenza vaccination responses in human beings have been analyzed computationally, but the results have not been consistent (2, 3, 7–10). One reason for these inconsistent results are the relatively small sample sizes. Another is that studies focus on only one biological aspect, for example molecular correlates of protection by using transcriptome data (11). However, a more robust approach to understanding how a vaccine works would involve analyzing multiple parameters from many individuals across different populations to more accurately capture biological variability. Furthermore, this would increase the statistical power, ultimately leading to the generation of classification and regression models with more robust performance metrics. While the number of studies and the amount of data are expanding dramatically, analyzing diverse samples across clinical studies remains challenging (12). This is particularly true for data from flow and mass cytometry where the number of markers analyzed can vary tremendously (13).

In this study, we develop an approach that optimizes a machine learning workflow through a Sequential Iterative Modeling “OverNight” (SIMON). SIMON is specifically tailored for clinical data containing inconsistent features with many missing values. SIMON utilizes multi-set intersections to successfully feed such data into an automated machine learning process with minimal sample losses. Our approach runs hundreds of different machine learning algorithms to find the ones which fit any given data distribution, and this maximizes predictive accuracy and other performance measurements. We used SIMON to analyze data from the Stanford Human Immune Monitoring Center (HIMC) collected from five separate clinical studies of seasonal influenza vaccination, obtained over eight years, with various platforms and expanding parameters. This enabled a systems-level identification of features that correlate with protective immunity to influenza. In the resulting models, we identified several previously unknown immune cell subsets that correlated with a successful influenza vaccination outcome, as defined by antibody responses. The impact of our findings is twofold. First, the study offers a new tool that can increase the accuracy of predictions from heterogeneous biological datasets. Second, it provides new targets for the development of the next-generation of influenza vaccines.

## Results

### Subhead 1. Preprocessing of data collected across different clinical studies

To test robustness of our approach, we used data from the Stanford HIMC. This data included 187 nominally healthy individuals between 8 and 40 years of age undergoing an annual influenza vaccination recruited over eight consecutive seasons, from 2007 to 2014, and five clinical studies (**Fig. 1A**). Blood samples were acquired before vaccination and on day 28 after vaccination. Over 3,800 parameters were measured at baseline. This included 102 blood-derived immune cell subsets analyzed by mass cytometry (**Fig. S1, Table S1**). It also included the signaling capacity of over 30 immune cells subsets stimulated with seven conditions, which were evaluated by measuring the phosphorylation of nine proteins (**Table S2**). Additionally, up to 50 serum analytes were evaluated using Luminex bead arrays (**Table S3**). On day 28 after vaccination, the serum titer of haemagglutinin-specific antibodies against all vaccine strains was determined using the hemagglutination inhibition assay (HAI), which is the best-defined correlate of influenza immunity induced by this vaccine (14). The HAI antibody titers were calculated as the fold change between the HAI titer at day 28 relative to the baseline titer. High and low responders were determined using metrics defined by the US Centers for Disease Control to evaluate influenza vaccine efficacy: seroconversion and seroprotection (15). Individuals were considered to be high responders if they had a protective HAI antibody titer to all vaccine strains (HAI antibody titer > 40) and if they seroconverted (geomean HAI titer > 4).

**Fig. 1.**
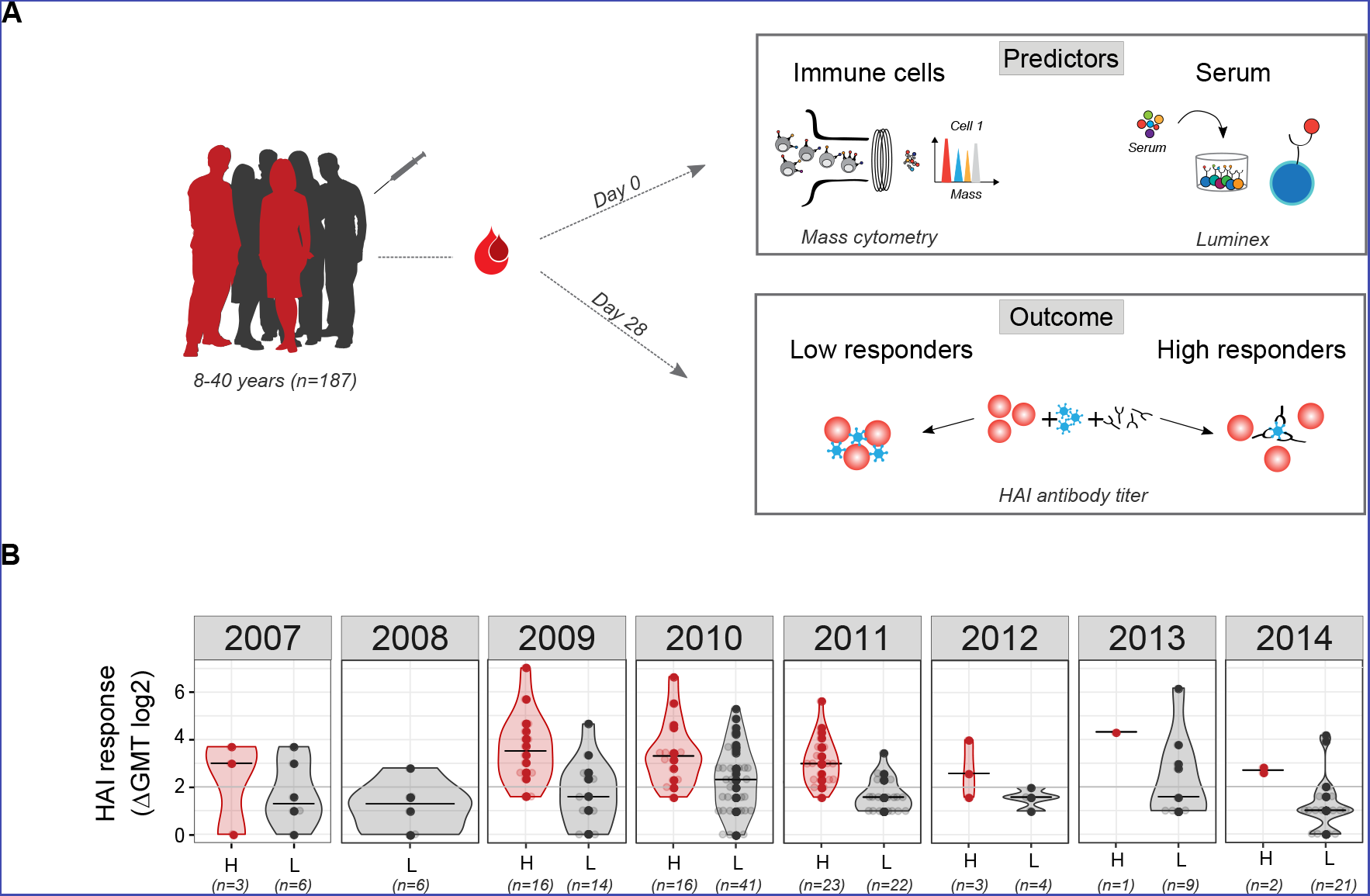
Study design. **(A)** One hundred and eighty-seven healthy donors (8-40 years of age) were recruited across eight consecutive influenza seasons. Data acquired at the baseline (day 0) included phenotypical and functional state (phosphorylated proteins) of immune cells analyzed using flow or mass cytometry and serum analysis using Luminex assay. Individuals were labelled as high or low responders, depending on the HAI antibody titers determined on day 28 after vaccination. **(B)** HAI antibody responses to influenza vaccine strains in high (H, red) and low (L, grey) responders across years. Numbers below x-axis indicate the number of donors in each group. HAI responses are shown as geometric mean titer (GMT) calculated as a fold change between day 0 and day 28 after vaccination for all vaccine strains. Violin plots show distribution of individuals. The line shows the median. Seroconversion is defined as 4-fold increase in HAI titer for all vaccine strains (denoted by a grey line).

Out of 187 analyzed donors, 64 were identified as high responders and 123 as low responders (**Fig. 1B**). Overall, there were no major differences in the age, gender, or study year between the high and low responders (**Fig. S2**). The only exception was that a higher proportion of adolescents were high responders, which is in line with published data(16) (**Fig. S2B**).

### Subhead 2. Dealing with missing values using intersection function

A major problem when using data across clinical studies and years is the lack of overlap between the features measured. Indeed, even though the data comes from a single facility, in many years there was an increase in the number of parameters measured, especially in the transition from FACS analysis (12-14 parameters) to mass cytometry (25-34 parameters). Since all assays were not performed across all studies and years (**Fig. S3**), our initial dataset had many missing values. The dataset contained 187 rows/donors and 3,284 columns/features, yielding a total of 614,108 values. However, 572,081 values were missing, resulting in high data sparsity. That is, the percentage of missing values in the dataset was 93.2% (**Fig. S4**). Such high data sparsity, which is commonly encountered in the clinical data, does not allow for straightforward statistical analysis. Therefore, we had to reduce the number of missing values. Researchers and data scientists deal with missing values either by deletion or by imputation of missing data (17). However, analysis of the missing data distribution revealed that when all studies were combined, the dataset had missing values in every column and every row and many of the columns and rows had sparsity of 90% (**Table S4**). Therefore, if we deleted either rows or columns, this would result in data with zero subjects. This approach was unsuitable. Additionally, effective imputation was strongly limited by the small number of cases that could be used as prior knowledge. Overall, we concluded that the high number of columns and rows with missing values made it impossible to use the whole dataset for further analysis.

Since this could be a very useful dataset for predictive modeling of influenza vaccine responses, we explored alternative ways to reduce the number of missing values. To ensure that interpretation of the initial dataset was preserved and so as not to introduce bias, we selected feature subsets from the original dataset without transformation by identification of the overlap (i.e. intersection) between multiple donors. We hypothesized that by using intersection, we could identify features shared across donors. Such a process could generate feature subsets that span an entire initial dataset. Additionally, it was expected that reducing the number of features would improve the performance of the model, such as was shown for random initial subset selection (18).

In the first step of SIMON, we implemented an algorithm, *mulset,* to identify features shared across donors and generate datasets containing all possible combinations of features and donors across the entire initial dataset (**Fig. 2**). While strategies to find an intersection among large number of sets have been reported(19), detecting intersections in a dataset with 614,108 datapoints would be challenging. The *mulset* was inspired by an approach commonly used in computer science to accelerate detection of duplicated records across large databases (20). By using the intersect function, we identified shared features between donors. These were converted to a unique shared feature identifier using the hash function. This process allowed the rapid identification of donors with shared features and the generation of datasets that can be used in further analysis. The *mulset* algorithm calculated overlapping features between all donors, resulting in 34 datasets with different numbers of donors and features (**Table S5**). After applying the *mulset* algorithm, the dimensionality of the data was significantly reduced, since all generated datasets had a maximum of 300 shared features. Eleven of the generated datasets had a higher number of donors than features, with a maximum number of 143 donors that shared 49 features (**Table S5, dataset 8**). Overall, the first step in the SIMON produced more restricted datasets with higher data quality and reduced the number of features, making possible to continue the data analysis.

**Fig. 2.**
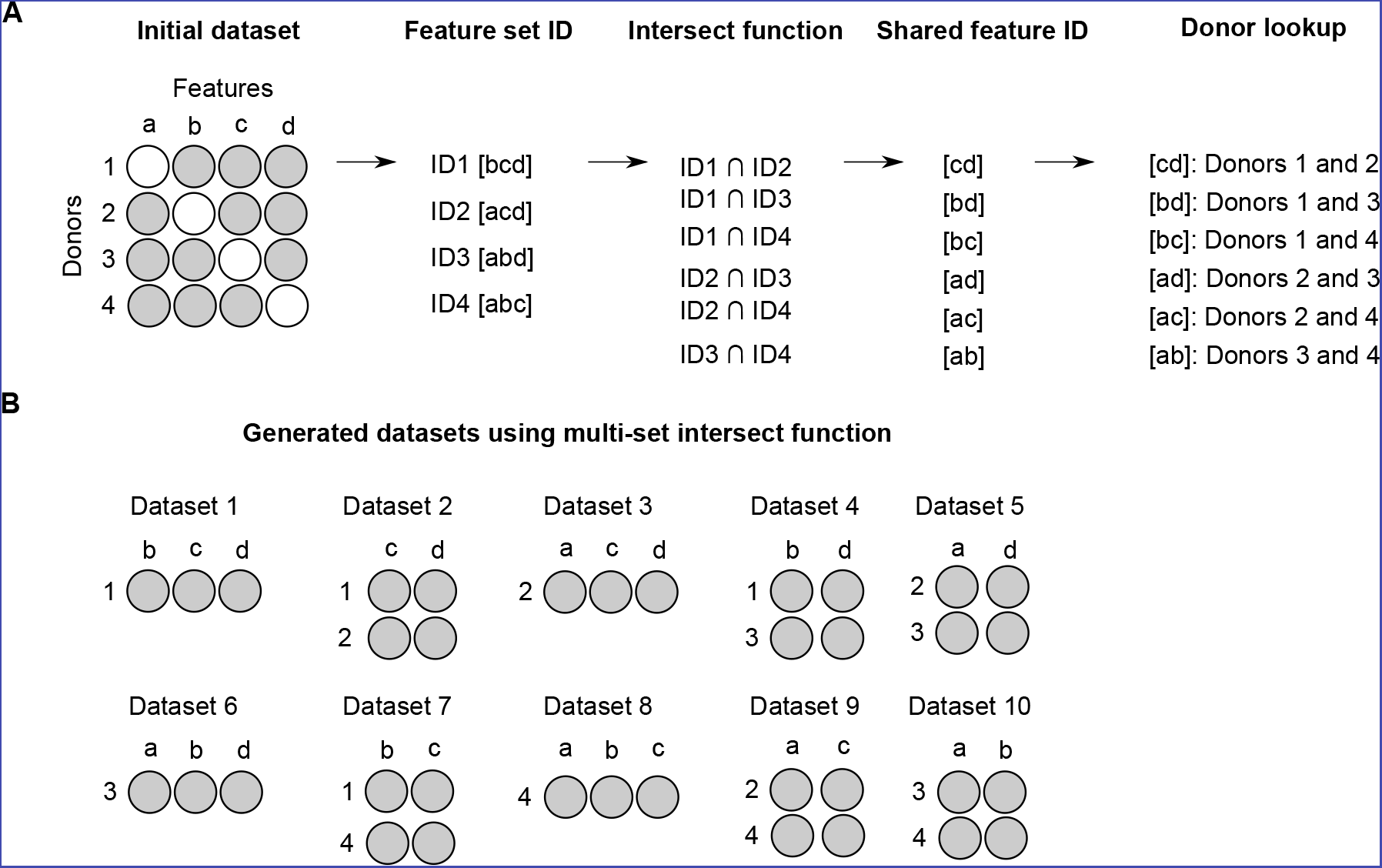
Automated feature subset generation using multi-set intersect function. Schematic example showing the initial dataset with four features and four donors. Missing values are indicated by white circles. Missing values are present is such a way that either removal of donors or features would result in no data for analysis. **(A)** Using a multi-set intersect function, the *mulset* algorithm, identified shared feature sets between donors. First, for each donor, the algorithm determined the unique feature ID. Second, using the intersect function, it identified shared features, which were then converted to shared features ID using hash functions. Finally, the *mulset* algorithm searched the database and identified donors with shared feature sets. **(B)** *Mulset* generated ten distinct datasets with defined feature and donor numbers, as indicated.

### Subhead 3. Automating the machine learning process and feature selection

The next step, following data preprocessing, was to apply machine learning algorithms to extract patterns and knowledge from each of the 34 datasets. To select relevant features, we based our approach on the method for feature selection proposed by Kohavi and John (21). In the original approach, termed wrapper, feature subsets were selected using two families of algorithms: the decision trees and the Naïve-Bayes (21). In this study, we build upon this approach by adding ensemble algorithms (of which Random Forest was previously shown to be suitable for feature selection (22)) and other dimensionality-reduction algorithms, such as discriminant analysis. It is widely recognized that a best algorithm for all datasets does not exist (23). Currently, choosing an appropriate algorithm is done through a trial-and-error approach, with only a few algorithms tested. To identify optimal algorithms more quickly and efficiently across a broad spectrum of possibilities, we implemented an automated machine learning process in SIMON.

SIMON is described briefly in **Fig. 3**. The feature subset selection was performed by testing multiple algorithms without any prior knowledge and user-defined parameters on each of the 34 datasets in a sequential and iterative manner. First, each dataset was split into 75% training and 25% test sets, preserving balanced distribution of high and low responders. The training set was used for model training and feature selection. The accuracy of the feature selection was determined using a10-fold cross-validation, which was shown to out-perform other resampling techniques for model selection (24). The test set was used for evaluating model performance on independent data not used in the model training. In general, it is most efficient to train the model on the entire dataset. However, in our case, it was important to have an independent test set to evaluate and then compare performance of the many models we expected to obtain. Additionally, evaluating model performance using only cross-validation is not sufficient to conclude that model can be applied to other datasets. There could be a problem with overfitting, such as when a model does not generalize well to unseen data. Second, a fully automated process of model training utilizing 128 machine learning algorithms was done initially on the training set and repeated for each dataset. **Table S6** provides a list of all machine learning classification algorithms used. Each model was evaluated by calculating the performance parameters using the confusion matrix on the training and test sets. A confusion matrix calculates false positive and negatives, as well as true positive and negatives. This allows for more detailed analysis than accuracy, which only gives information about the proportion of correct classifications, and therefore can lead to misleading results (25). In SIMON, for each model we calculated the proportion of actual positive cases that were correctly identified (i.e. sensitivity) and the proportion of correctly identified actual negative cases (i.e. specificity). All performance parameters were saved in the MySQL database. Finally, to compare the models and discover which performed best, we calculated an Area Under the ROC Curve (AUROC). This is a widely-used measure of quality for the classification of models, especially in biology (26). A random classifier that cannot distinguish between two groups has AUROC of 0.5, while AUROC for a perfect classifier that separates two groups without any overlap is equal to 1.0 (27). Therefore, the training and test AUROC are reported throughout the text and models are compared using that metric of performance.

**Fig. 3.**
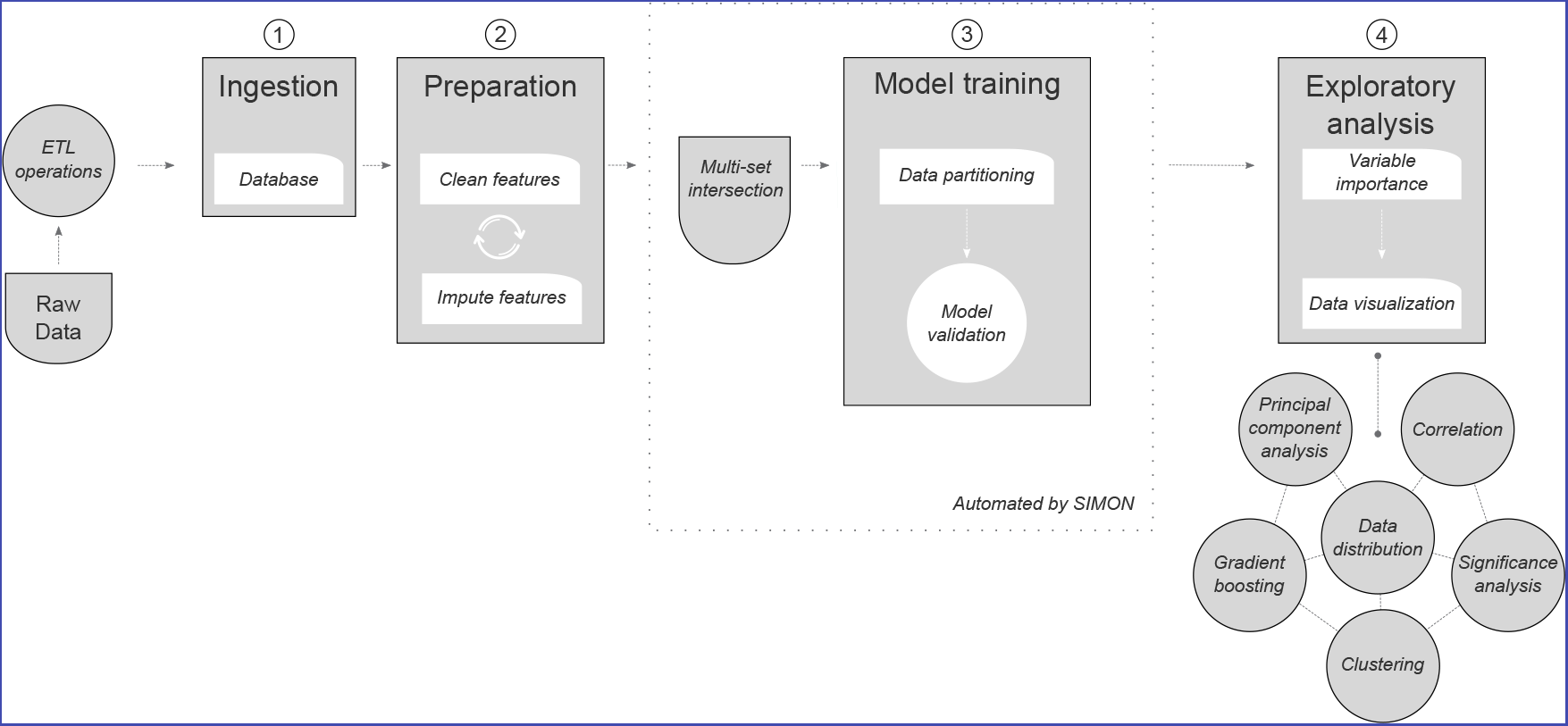
Automated feature selection and machine learning process integrated in SIMON. Before building a model, raw data were processed (cleaned, corrected, normalized, etc.) using extract-transform-load (ETL) operations and the database was built. In the second step, new features were created from the existing data, GMT of the HAI response was calculated, and individuals were labelled as high or low responders. Third, datasets were generated using multi-set intersection function. Each dataset was then used for model training in a fully automated machine learning process, implemented in SIMON. Briefly, before training started, each dataset was partitioned into training and test sets, which were excluded from the model-building phase. Finally, in the exploratory analysis, each model was evaluated based on its performance, and features were selected based on the importance score.

To test the feasibility of SIMON, we ran more than 2,400 machine learning analyses on 34 datasets. SIMON built models for 19 datasets, with an average of 54 models built per dataset (**Table S7**). None of the 128 machine learning algorithms tested were able to build a model for 15 of the datasets. This indicates that those have poor data quality and distributions. Therefore, they were discarded from further analysis. With the remaining 19 datasets, models were built with the training AUROC values ranging from a minimum of 0.08 to a maximum of 0.92 (**Table S7**). Overall, the automated machine learning process improved the performance of the models in all 19 datasets, with a gain of performance ranging from 30 to 91% (**Table S7**). This indicates that SIMON facilitates the identification of optimal algorithms, which ultimately increases the performance of models.

### Subhead 4. Performance estimation and model selection

Before model comparison, other performance parameters were calculated, in addition to AUROC, and were used to filter out poorly performing models with the goal of facilitating further exploratory analysis. To remove random classifiers, all models with AUROC ≤ 0.5 on both training and test sets were discarded. Furthermore, all models in which specificity and sensitivity of both training and test sets were < 0.5 (i.e. models with higher proportion of false positive and negative values) were also removed. This restriction discarded models in which the classifier achieved high performance, as indicated by a high AUROC, at the cost of a high false positive or negative rate (28, 29). After applying these filters, many models were removed, decreasing the average number of models per dataset to three (**Table S8**). Additionally, eight datasets were discarded. This filtering step was essential to remove models which would otherwise be falsely evaluated as high performing, such as those built using dataset 205, for which a high AUROC of 0.92 was obtained at the expense of low specificity (0.06) (**Table S7**).

To compare models within one dataset and discover which performs best, the random number seed was set before training with each algorithm. This ensured that each algorithm trained the model on the same data partitions and repeats. Further, it allowed for comparison of models using AUROC. In general, AUROC values between 0.9-1 are considered excellent, 0.8-0.9 good, 0.7-0.8 fair and values between 0.6-0.7 are considered as having poor discriminative ability (30). In SIMON, models trained on six datasets were built with fair discriminative ability (max. train AUROC between 0.7-0.8) (**Table S8**). To avoid overfitting, we additionally evaluated the performance of each model on the test set, which was not used for building the model. In this case, models trained on the three datasets were built with a fair discriminative ability (**Table S9, datasets 5, 13, and 171**). One dataset (**Table S9, dataset 36**) was built with a good discriminative ability (max. test AUROC 0.86), which could be generalized to an independent set. It should be noted that maximum AUROC values did not necessarily come from the same model (e.g. maximum train AUROC might come from model 1, while maximum test AUROC from model 2). To account for that, we add another filter to remove all models with poor discriminative ability, that is all models in which the train and test AUROC were less than 0.7. By applying this restriction, we were left with only two datasets (datasets 13 and 36). These were used for further analysis and feature selection. The model build on dataset 36, with the shrinkage discriminant analysis, out-performed the other four models as evaluated by comparison of train AUROC (**Fig. S5A, Table S10**). A model was built with train AUROC of 0.78 and it performed well on an independent test set (test AUROC 0.86). The model build on dataset 13 with the Naïve-Bayes performed better than the other model built for the same dataset (train AUROC 0.75, test AUROC 0.7) (**Fig. S5B, Table S11**).

Overall, SIMON facilitated exploratory analysis and discovery of models with good discriminative performance by integrating the filtering steps and evaluating comprehensive model performance.

### Subhead 5. Identification of all-relevant cellular predictors using SIMON

After selection of the best-performing models, we focused on feature selection. Our goal was to use SIMON to identify all-relevant features to deepen our knowledge about the process that drives antibody generation in response to influenza vaccination. To solve this problem, classifiers were used in SIMON to rank features based on their contribution to the model. Features were ranked depending on the variable importance score calculated for each model (31). The score ranges from 0 to 100. Features with variable importance score of 0 are not important for the classification model and can be removed from training the model.

First, we focused on the model built on dataset 13 with 61 donors (**Table S12**). Out of 76 features, 64 had measurable variable importance score and 15 features had variable importance score above 50 (**Fig. 4A**, **Table S13**). The top**-**ranked feature that highly contributed to this model was CD4^+^ T cells with the CD127^-^CD25^hi^ phenotype (described as regulatory T cells or Tregs(32)) that expressed CD161 and CD45RA markers (**Table S13, rank 1**). The frequency of Tregs with CD161^-^CD45RA^+^ phenotype was shown to be significantly greater among the high responders (**Fig. 4B**, FDR < 0.01). To further explain features that contributed to this model, we performed correlation analysis. Correlation analysis revealed that Tregs with CD161^-^CD45RA^+^ phenotype had a significant positive correlation with the top-ranked feature, CD161^+^CD45RA^+^ Tregs (Pearson’s r = 0.54, p < 0.0001 after multiple comparison adjustment using the B-H correction) (**Fig. S6**). Additionally, CD161^+^CD45RA^+^ Tregs had a weak, but significant positive correlation with CD161^+^ CD4^+^ T cells (Pearson’s r = 0.08, p = 0.001 after multiple comparison adjustment using the B-H correction), which had high variable importance score (**Table S13, rank 9**). Such correlation indicated that these subsets might describe similar family of CD4+ T cells contributing to the generation of antibody responses after influenza vaccination. Indeed, a recent study suggests that expression of CD161 marks a distinct family of human T cells with a distinct lineage and with innate-like capabilities (33).

**Fig. 4.**
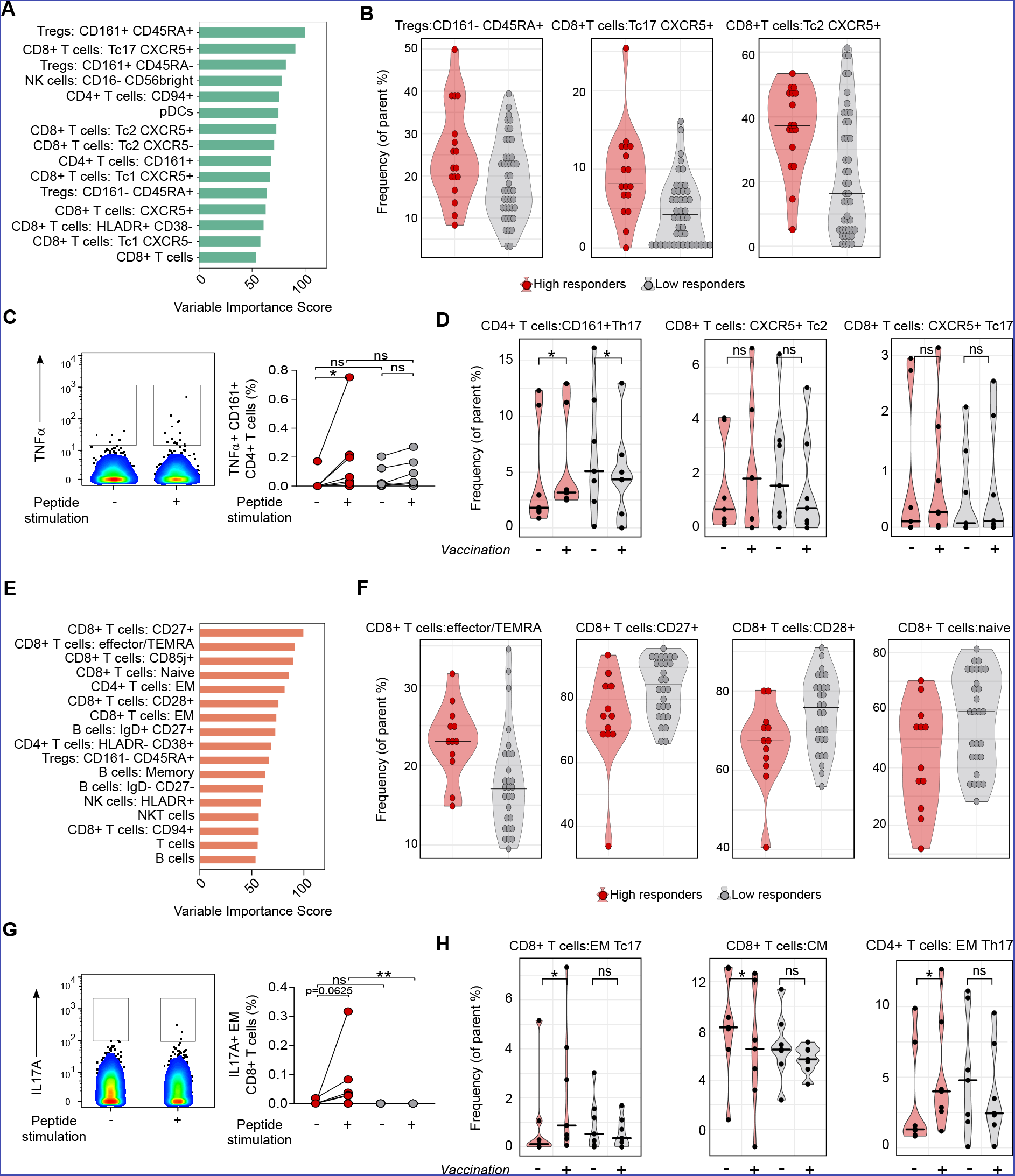
SIMON identifies cellular signature associated with the successful generation of influenza immunity after vaccination. **(A)** Features with variable importance score above 50 from the model built on dataset 13 are shown. **(B)** Raw data confirmed by SAM analysis to be significantly changed in the donors from dataset 13, indicating frequency of cells (as a percentage of the parent population). **(C)** Representative plot showing TNFα intracellular staining of CD161^+^ CD4^+^ T cells in the unstimulated (-) or influenza peptide pool (+) stimulated PBMC from high responder obtained before vaccination. Graph on the right shows the frequency of TNFα^+^ CD161^+^ CD4^+^ T cells from high responders (red circles) and low responders (grey circles) in the samples before vaccination. Individual donors are connected with lines. **(D)** Violin plots show distribution of frequency of CD161^+^ CD4^+^ T cells and CXCR5^+^ CD8^+^ T with Tc2 and Tc17 phenotype in the PBMC samples derived from high (red, n = 7) and low responders (grey, n = 7) analyzed before vaccination (-) and on day 28 after vaccination (+). **(E)** Variable importance score of features selected in the model built on dataset 36 with score above 50. **(F)** Significant immune cell subsets selected by SAM analysis shown as raw data corresponding to donors from dataset 36, indicating frequency of cells (as percentage of parent population). **(G)** Representative plot showing IL17A intracellular staining of EM CD8+ T cells in the unstimulated (-) or influenza peptide pool stimulated (+) PBMC from high responders, obtained after vaccination. The graph on the right shows the frequency of IL17A+ EM CD8+ T cells from high (red circles) and low (grey circles) responders in the samples after vaccination. **(H)** Violin plots show distribution of frequency of CD4^+^ and CD8^+^ T cells, with indicated phenotypes analyzed in the PBMC samples derived from high (red, n = 7) and low responders (grey, n = 7) before (-) and on day 28 after (+) vaccination. Graphs shown in **(C, D, G and H)** represent combined data from seven independent experiments. Violin plots show distribution of individuals. These are represented by red circles for high responders and grey circles for low responders. The line indicates the median. Statistical analysis between high and low responders was performed with one-way ANOVA Kruskal-Wallis test followed by Dunn’s multiple comparison test. Analysis within groups before and after vaccination was calculated using two-tailed Wilcoxon matched-pairs signed rank test. Significance in SAM analysis was considered at FDR < 0.01. ns - not significant, *p < 0.05, **p < 0.01.

To experimentally validate results from this model, we analyzed the phenotype and functionality of immune cells before and after vaccination in the independent samples from 14 individuals (7 high and 7 low responders) (**Table S14**). We found that after stimulation with the influenza peptides, CD161^+^ CD4^+^ T cells from high, but not low responders, produced TNFα in the samples prior to vaccination (**Fig. 4C**). This indicated that CD161^+^ CD4^+^ T cells from high responders had a pool of pre-existing influenza-specific T cells. Additionally, after vaccination, the frequency of CD161^+^ CD4^+^ T cells with a CCR6^+^ CXCR3^-^ (Th17) phenotype in high responders increased significantly (**Fig. 4D**).

The second most important feature in this model was CXCR5^+^ CD8^+^ T cells (also known as follicular cytotoxic T cells) (34–36) with a CCR6^+^ CXCR3^-^ (Tc17) phenotype (**Table S13, rank 2**). Frequencies of CXCR5^+^ CD8^+^ T cells with Tc17 were significantly increased among the high responders (**Fig. 4B**, FDR < 0.01). Additionally, frequencies of CXCR5^+^ CD8^+^ T cells with a CCR6^-^CXCR3^-^ (Tc2) phenotype were also increased in the same group (**Fig. 4B**, FDR < 0.01). CXCR5^+^ CD8^+^ T cells with Tc2 phenotype were also identified as important in this model (**Table S13, rank 7**) and had a significant positive correlation with Tc17 CXCR5^+^ CD8^+^ T cells (Pearson’s r = 0.66, p < 0.0001 after multiple comparison adjustment using the B-H correction) (**Fig. S6**). However, analysis of the experimental data showed no significant participation of CXCR5^+^ CD8^+^ T cells in vaccine-induced responses, even though in a few of the high responders there was an increase of CXCR5^+^ CD8^+^ T cells with a Tc2 and Tc17 phenotype (**Fig. 4D**).

The results obtained in this model were confirmed using an R package, Boruta, that implements a novel feature selection algorithm for identifying all relevant features (22). CD127^-^CD25^hi^ CD4^+^ T cells with the CD161 expression and CXCR5^+^ CD8^+^ T cells with Tc2 or Tc17 phenotype were identified as important (p < 0.05, after multiple comparison adjustment using the Bonferroni method), confirming findings obtained by SIMON (**Fig. S7A**).

Second, we explored the features selected in the better performing model built on dataset 36, comprising of 40 donors (**Table S15**). Out of 103 features, 88 had measurable variable importance scores ranging from 5 to 100 (**Table S16**). Of those, 17 features had a variable importance score above 50 (**Fig. 4E**), indicating a strong contribution for this classification model. Interestingly, the effector memory (EM) CD4^+^ T cells, previously reported to correlate with antibody responses to influenza vaccine (37), were ranked in 5^th^ place in our model. Moreover, B cells with memory phenotype, including a subset of IgD^+^ CD27^+^ memory B cells identified in previous studies (3, 8, 38), contributed to our model (**Table S16**). Obtaining results supported by other studies gave us confidence in further analysis of our classification model. Importantly the top four features identified have not previously been implicated as playing a major role in antibody responses to influenza vaccination, or indeed any antibody response. These included CD8^+^ T cells with expression of CD27 or CD85j markers, and CD8^+^ T cells with varying degree of expression of CCR7 and CD45RA markers, described as naïve, effector or terminally differentiated effector (TEMRA) and memory subsets (39). Analysis of the data particularly indicated that effector/TEMRA CD8^+^ T cells increased significantly among high responders (**Fig. 4F**, FDR < 0.01). In contrast, low responders had significantly higher frequency of early CD27^+^/CD28^+^ CD8^+^ T cells and naïve CD8^+^ T cells (**Fig. 4F**, FDR < 0.01). Moreover, the effector/TEMRA CD8^+^ T cells were confirmed to contribute to this model by Boruta (p < 0.05, after multiple comparison adjustment using the Bonferroni method) (**Fig. S7B**).

The top four features that contributed the most to this model, were CD8^+^ T cells in early or late effector or memory states, indicating they might all be contributing to the influenza response through the same underlying mechanism. Indeed, correlation analysis showed that the top ranked subset, CD27^+^ CD8^+^ T cells, had a significant correlation coefficient with other subsets (naïve CD8^+^ T cells r= 0.80, CD28^+^ CD8^+^ T cells r = 0.85, CD85j^+^ CD8^+^ T cells r = −0.69, effector/TEMRA CD8^+^ T cells r = −0.61 and EM CD8^+^ T cells r = −0.71, p < 0.0001 after multiple comparison adjustment using the B-H correction) (**Fig. S8**). Additionally, a specific subset of CD8^+^ T cells expressing NK-cell-related receptor CD85j was identified as the TEMRA subset (40), while the expression of CD27 or CD28 was indicative of the subsets of T cells with a naïve or early differentiation phenotype (41).

In the analysis of the independent samples, EM CD8+ T cells from high responders produced IL17A after influenza peptide stimulation, demonstrating that this population contained influenza-specific T cells (**Fig. 4G**). Furthermore, the frequency of EM CD8^+^ T cells with a Tc17 phenotype was significantly increased only in high responders after vaccination (**Fig. 4H**). Additionally, the frequency of EM CD4+ T cells with Th17 phenotype was also increased in the same group of high responders after vaccination (**Fig. 4H**).

In summary, SIMON allowed us to identify both known and novel immune cell subsets that correlate with a robust antibody response to seasonal influenza vaccines. Particularly surprising were the number of different CD8^+^ T cell subsets, which are not typically thought of as playing any role in promoting robust antibody responses. We confirmed that IL17A producing EM CD8^+^ T cells, which contained a pool of pre-existing influenza T cells, were elevated in the high versus low responders with independent samples.

## Discussion

In this study, we developed a novel computational approach, SIMON, for the analysis of heterogenous data collected across years and from heterogenous datasets. SIMON increases the overall accuracy of predictive models by utilizing an automated machine learning process and feature selection. Using the results obtained by SIMON, we identified previously unrecognized CD4^+^ and CD8^+^ T cell subsets associated with robust antibody responses to seasonal influenza vaccines.

The accuracy of the machine learning models presented in this work was improved in two stages. First, to interrogate the entire dataset across different clinical studies, we integrated into SIMON an algorithm, *mulset*, which generates datasets using multi-set intersections. This is particularly suitable for data with many missing values. In our case, due to the high sparsity of initial dataset, this step was essential for the further analysis. In general, clinical datasets are often faced with the same problem, namely, that many features are measured on a small number of donors. Due to the rapid advance of immune monitoring technology, many more parameters in our studies were measured in the later years than earlier. The same situation might arise when combining data collected in different facilities. An alternative approach might be the imputation of the missing values, but this would likely introduce bias. Moreover, the major limitation of effective imputation is the number of cases that could be used as prior knowledge. The sparsity of our initial dataset was too high for effective imputation. By using intersections, SIMON selects feature subsets by preserving the interpretation of the initial dataset and without introduction of a bias. Overall, an automated feature intersection process increases statistical power by accounting for variability among different individuals. Potentially, it could be applied across clinical studies. Additionally, by reducing the number of features, this process improves the performance of models. This will be particularly important for the application of SIMON on larger publicly available datasets such as those stored in Gene Expression Omnibus repository (42) or ImmPort (43).

Second, finding the machine learning algorithm most suitable for specific data distribution allows for a better understanding of the data and provides much higher accuracy. The current state-of-the-art in building predictive models is to test several machine learning algorithms to find the optimal one. However, a single algorithm that fits all datasets doesn’t exist. If an algorithm performs well on a certain dataset, it doesn’t necessarily translate well to another dataset (even if it pertains to a closely related problem) (23). The overall accuracy of the predictive models depends on rigorous algorithm selection. With so many machine learning algorithms available, choosing the optimal one is a time-consuming task, often performed in a limited way (only dozens of algorithms are tested). Recent work has shown that automated machine learning can identify optimal algorithms more quickly and efficiently (44–46). Open competitions and crowdsourcing (e.g. www.kaggle.com), in which many groups contribute machine learning algorithms to build models for the same datasets, increases the accuracy and predictive performance of the models (47). By developing an automated machine learning process in SIMON, we can quickly identify the most appropriate machine learning algorithm (of the 128 tested) for any given dataset. Additionally, SIMON offers an alternative perspective on the application of algorithms that might never be used due to lack of expertise or knowledge necessary for their implementation. These features of SIMON also allow biologists with domain knowledge but who are not computationally adept to find the most effective tools with which to analyze their data.

In this study, we demonstrate the utility of SIMON and its automated machine learning processes to discover the principal features that correlate with high versus low influenza vaccine responders. We found it to be essential for identifying the best-performing models and extracting the most important features that contribute to those models. Performance of each model built in SIMON was automatically evaluated on both training and left-out test sets using well-known measures, such as AUROC, specificity, and sensitivity. This ensured that the model was not overfitted and that it could generalize to unseen data. Both models were selected by stringent restrictions in the exploratory analysis and were built with AUROC scores between 0.7 - 0.8. Nevertheless, since the goal of the study was to identify features that discriminate between high and low responders in a high-throughput manner, these models were built using the algorithms without any user-defined parameters. Therefore, each model could be fine-tuned, and its predictive performance might be increased. This could be of interest for researchers interested in building predictive models to identify features for use in diagnostic tests. In the future, we plan to improve SIMON by implementing an automated tuning process for each model.

This study demonstrated the advantage of SIMON over the conventional approach, in which one machine learning program is chosen by successfully identifying the immune signature driving influenza immunity. Some of our findings, such as the importance of effector memory CD4^+^ T cells and subsets of memory B cells, had been identified in previous studies (2, 8, 9) serving to validate our approach. Additionally, SIMON has identified previously unappreciated T cell subsets that discriminate between high and low responders. It is well known that T cells, in contrast to antibodies produced by cells of B lineage, have the ability to provide durable and cross-protective immunity by targeting internal conserved viral epitopes (48, 49). Therefore, the CD4^+^ and CD8^+^ T cell subsets identified in this study could be useful targets for the development of broadly protective influenza vaccines. Influenza-specific CD4^+^ T cells have already been shown to be important for the generation of influenza immunity (50, 51). This was confirmed in the current study by showing that high responders had a pre-existing pool of influenza-specific CD4^+^ T cells expressing CD161. Additionally, we found that CD8^+^ T cells with an effector/TEMRA, EM and Tc17 phenotype and CXCR5 expression correlated with improved vaccine responses. These subsets are particularly interesting candidates and it will be of considerable interest to understand how they contribute to more robust antibody responses. CXCR5^+^ CD8^+^ T cells are enriched in the B cell follicles of germinal centers (35, 52) and they can promote B cell survival and antibody generation (36). CD8^+^ T cells with a Tc17 phenotype have been detected in the lungs of mice challenged with influenza A virus (53). Using independent samples from donors that weren’t included in the building and testing of our model, we found that CD8^+^ T cells from high responders contained influenza-specific cells with the ability to produce IL17A in response to peptide stimulation. In a mouse model, IL17A has been shown to be important for the generation of the antibody responses necessary to clear an influenza virus infection (54). This apparent role of IL17A in the modulation of antibody responses and proper functioning of germinal centers has only recently been described (55). Interestingly, CD161^+^ CD45RA^+^ Tregs, the other subset we identified, have also been described as memory cells with the ability to produce IL17A (56). Therefore, both cell types may provide IL17A.

Here, we demonstrate that a combination of systems biology tools, advances in the field of machine learning, and experimental investigation, provides a new and more efficient way to gain biological insight from complex datasets, despite high sparsity.

## Materials and Methods

### Subjects, sample and data collection

All clinical studies were approved by the Stanford Institutional Review Board and performed in accordance with guidelines on human cell research. Peripheral blood samples were obtained at the Clinical and Translational Research Unit at Stanford University after written informed consent/assent was obtained from participants. Samples were processed and cryopreserved by the Stanford HIMC BioBank according to the standard operating protocols (57). All materials and data were analyzed anonymously.

In this study, we used data from one hundred and eighty-seven healthy donors that were enrolled in influenza vaccine studies at the Stanford-LPCH Vaccine Program from the 2007 to 2014. This included the following studies: SLVP015 (NCT01827462, NIAID ImmPort accession number SDY212, data analysis described in (58)), SLVP017 (NCT02133781, NCT03020498, NCT03020537), SLVP018 (NCT01987349, NCT03022396, NCT03022422, NCT03022435, NCT3023176, data analysis published in (59)), SLVP021 (NCT02141581), SLVP028 (NCT03088904) and SLVP029 (NCT03028974). Individuals were selected for this study based on the following criteria: (1) age range from 8-40 years, (2) received inactivated influenza vaccine (IIV, Fluzone, intramuscularly), (3) only data from the first visit (some donors came in consecutive years), (4) HAI titer measured and (5) information about gender and age available. Exclusion/inclusion criteria, samples that were acquired with timepoints and analyses performed are described in the study record details at website repository for clinical studies (www.ClinicalTrials.gov) using provided identifiers. All the protocols for sample analysis such as immunophenotyping and determination of signaling responses to stimulation using flow or mass cytometry, HAI titer determination and determination of cytokines/chemokines in samples using Luminex assay are available online (57). Additionally, immunophenotyping using mass cytometry was published in Leipold and Maecker (60). Phosphoflow assay using flow cytometry (for studies SLVP15, SLVP18 and SLVP21 from 2007 to 2011), was described in (58, 59) or for mass cytometry (for study SLVP21 in 2013) (61). Luminex assay was described in (58, 59). The HAI assay was performed on sera from day 0 and day 28 using a well-established method (62) and was described before (2, 58).

All data used were analyzed and processed at the HIMC, as previously described (63) and uploaded to the Stanford Data Miner (64). Briefly, data from both Luminex assays were normalized at the plate level to mitigate batch and plate effects. The two median fluorescence intensity (MFI) values for each sample for each analyte were averaged, and then log-base 2 transformed. Z-scores ((value–mean)/standard deviation) were computed, with means and standard deviations computed for each analyte for each plate. Thus, units of measurement were Zlog2 for serum Luminex. For phospho-flow data acquired on flow cytometer a fold change value was computed as the stimulated readout divided by the unstimulated readout (e.g. 90th percentile of MFI of CD4+ pSTAT5 IFNα stimulated / 90th percentile of CD4+ pSTAT5 unstimulated cells), while for data acquired using mass cytometry a fold change was calculated by subtracting the arcsinh (intensity) between stimulated and unstimulated (arsinh stim – arcsing unstim). For immunophenotyping using mass cytometer units of measurement were percentage of parent population.

#### Aggregation of data and generation of feature subsets

The preprocessed data from Stanford influenza datasets were obtained from HIMC Stanford Data Miner (64). This included total of 177 csv files, which were automatically imported to the MySQL database to facilitate further analysis. Datasets were merged using shared variables, such as Donor ID, Study ID, gender, age, race, Donor visit ID, Visit year, Experimental data (connected to Donor visit ID), Assay, Name and value of the measured analyte. Data harmonization across different clinical studies was accomplished by introduction of feature termed Visit internal ID, which allowed us to discriminate between different visits of the unique donor in different years. We standardized names of the vaccines, for example TIV and IIV3 were named Inactivated influenza vaccine. Finally, we calculated the vaccine outcome parameter using HAI antibody titers. High responders were determined as individuals that have HAI antibody titer for all vaccine strains above 40 and geometric mean (GeoMean) HAI fold change >4. The fold change is calculated as: GeoMean HAI antibody titer for all vaccine strains on day 28 / GeoMean HAI antibody titer for all vaccine strains on day 0. To facilitate analysis, vaccine outcome was expressed as binary value: high responders were given value of 1, while low responders value zero.

The aggregated dataset contained 187 donors and 3,284 features, yielding a total of 614,108 datapoints. The dataset had 93.2% of missing values (572,081 missing values). To deal with missing values, in the first step of the SIMON we implemented a novel algorithm, *mulset* that allows for faster generation of datasets with all possible combinations of features and donors across initial dataset. To efficiently compute shared features and find quickly similarity between donors, *mulset* algorithm generated unique feature identifier for each donor. Then, intersection between the identifiers was used to identify shared variables. The identified, shared variables are then converted to unique shared features identifiers using hash function. Finally, data was exported from the database according to the shared features. In total, *mulset* generated 45 different datasets. To avoid proceeding with machine learning process using datasets with misleading results, we removed datasets with less than 5 features and less than 15 donors. After applying that restriction, 11 datasets were deleted, and final analysis was performed on 34 datasets.

### Overview of SIMON - Sequential Iterative Modeling “Overnight”

To identify baseline immune predictors that can discriminate between high and low responders following influenza vaccination, we applied sequential, iterative modeling “overnight”, shortly SIMON. The SIMON allows for dataset generation, feature subset selection, classification, evaluation of the classification performance and determination of feature importance in the selected models. The SIMON was implemented in R programming language (65) (**Data file S1**). First in SIMON we automated the process of dataset generation using *mulset* algorithm as described above. Next, each dataset was partitioned into 75% training and 25% test set with balanced class distribution of high and low responders using the function *createDataPartition* from the Caret package (31). To prevent evaluation of small test sets that would lead to misleading performance parameters, datasets with less than 10 donors in test sets were discarded. Next, the model training using 128 machine learning algorithms suitable for classification training (**Table S6**) was initiated for each train dataset. Test sets were hold-out for evaluation of model performance on unseen datasets. This step was crucial to prevent overfitting. All algorithms were processed in an automated way through the Caret library (31). Each model was evaluated using 10-fold cross-validation (24) repeated 3-times. Additionally, performance of each model was evaluated on the test set which was held out before model training by calculating performance from a confusion matrix using available R package (66). Furthermore, contribution of each feature to the trained model was evaluated and variable importance score is calculated as described (31). All prediction metrics and performance variables are stored in the MySQL database for the final exploratory analysis. Detailed description of the overall processes is as follows.

#### Model training and performance evaluation

For each dataset, model training was performed on train set using 128 machine learning algorithms (**Table S6**). All algorithms were implemented without any user-defined parameters and with the default tuning parameters, as described in the Caret library (31). For each model we defined type of resampling. In this study10-fold cross-validation repeated three times was used. Each model was then evaluated by calculating performance measures using the confusion matrix. Confusion matrix or contingency table is used to evaluate the performance of a classification model on a set of data for which the true values are known. The confusion matrix has four categories (see illustration below).

|  |  | **Actual** |  |
| --- | --- | --- | --- |
|  |  | Positive<br>(High responder) | Negative<br>(Low responder) |
| **Predicted** | Positive<br>(High responder) | True positives (TP) | False negatives (FN) |
|  | Negative<br>(Low responder) | False positives (FP) | True negatives (TN) |

True positives (TP) are cases in which classification model predicted they are high responders and indeed those cases were high responders, while true negatives (TN) correspond to cases correctly labeled as low responders. Finally, false negatives (FN) and false positives (FP) refer to low responders or high responders that were incorrectly labeled. From a confusion matrix, to evaluate classification models we calculated following performance measures. Accuracy, a measure how often the classifier is correct was calculated as (TP+TN)/total number of observations. Specificity, the proportion of actual negative cases (low responders) that were correctly identified was calculated as TN/(FP+TN), while sensitivity (also known as recall or true positive rate), the proportion of actual positive cases (high responders) correctly labeled was calculated as TP/(TP+FN). To summarize the performance of classification models over all possible thresholds, we generated the Receiver Operating Characteristic (ROC) curve by plotting the sensitivity (y-axis) and the false positive rate (the proportion of low responders misclassified as high responders), which was calculated as 1-specificity (x-axis). Finally, we calculated the are under the ROC curve (AUROC) using an R package (66) and used this measure to summarize the performance of the models. AUROC has values between zero and one, and higher values indicate better performance. Value of 0.5 indicates a random classifier, and this was used as a cutoff to remove classifiers that could not distinguish between high and low responders better than by random chance. In this study, 10-fold cross validation was applied three times, the AUROC was calculated for each repeated iteration, and the average AUROC (and other measures) are reported as an overall quantitative estimate of classification performance. Additionally, before model training, same seed for random number generator was applied (*set.seed* 1234). This resulted in the uniformity where for each model same resamples were used for performance evaluation. From this, we compared models and evaluated which model was performing better in terms of AUROC values by comparing performance of the resampling distributions using functions described in the Caret (31).

#### Independent evaluation of the trained model

The performance of each model was additionally evaluated on the test set which was held out before training the model (25% of the dataset). The performance on the test set was evaluated exactly as described for the train set above. Confusion matrix was built and all the performance measures, including the AUROC, as for train set were computed. Test AUROC was used to select models, in addition to train AUROC.

#### Variable importance score

Contribution of each feature to the model i.e. variable importance score was calculated using the Caret library (31). Briefly, evaluation of the variable importance was calculated directly from the model specific metrics and the variable importance scores were scaled to have a maximum value of 100. Since in SIMON we utilized many different algorithms, the contribution of each feature to the model was estimated using the methods appropriate for each algorithm, as described in R packages (see reference list for the **Table S6**).

### Feature selection using Boruta algorithm

To evaluate the all-relevant features for the selected top-performing models built on datasets 13 and 36, we used an R package Boruta (22). Boruta algorithm performs as a wrapper algorithm around Random Forest (22). The method is suitable for selection of all-relevant features, and this is accomplished by comparing original features’ importance with importance achievable at random (estimated using permuted copies of the original features, called shadow features). In each iteration, Boruta removes irrelevant features and evaluates the performance of the model. Finally, analysis is finished either when all features are confirmed or rejected or when Boruta reaches a specified limit of runs. Boruta was performed using following parameters: maximal number of importance source runs, *maxRuns* at 1000, *pValue* confidence level 0.05, a multiple comparisons adjustment using Bonferroni method was applied (*mcAdj* set to TRUE), feature importance was obtained using Random Ferns (function *getImpFerns*) and to ensure reproducibility of the results we set the seed for the random number generator (*set.seed* 1337). Tentative features were also included returned in the Boruta results (*withTentative* argument was set to TRUE).

### Peptide stimulation and intracellular cytokine staining using mass cytometry

Thawed PBMC were rested in X-VIVO™ 15 medium (Lonza) supplemented with 10% FCS and human serum AB (Sigma) for 2 days at 10^7^ cells/ml in 24-well plate following “RESTORE” protocol (67, 68). For stimulation assay, 5×10^6^ PBMC were seeded in 96-well V-bottom plates (10^6^ PBMC/well) and stimulated overnight (12-16h) with the influenza overlapping peptide pool. Influenza peptide pool contained 483 peptides (20mers with 11aa overlap, Sigma Aldrich) spanning the entire influenza proteome from the influenza strain A/California/07/2009 (dissolved in DMSO at 20mg/ml, working concentration 0.2ug/ml/peptide) and 24 peptides with HLA-A*0201-specificity (9-10mers, Sigma Aldrich) generated against influenza proteins (hemagglutinin, nucleocapsid protein, matrix protein 1, nonstructural protein 1 and 2) from the influenza strain A/California/07/2009 using prediction software NetCTL-1.2 (69) (dissolved in water or PBS/DMSO at 20mg/ml, working concentration 2ug/ml/peptide) (**Table S18**). In both assay unstimulated sample was prepared in which only medium without peptides containing 0.5% DMSO was added. Protein transport inhibitor cocktail (eBioscience/Thermo Fisher) and antibody against CD107a were added at the beginning of the assay. After peptide stimulations, PBMC were washed with the CyFACS buffer (PBS supplemented with 2% BSA, 2 mM EDTA, and 0.1% soium azide) and stained with surface antibody cocktail (**Table S17**), filtered through 0.1um spin filter with 20uL/sample of Fc block (ThermoFisher) for 30min at 4°C. Cells were then washed with CyFACS buffer and incubated for 5min at RT in 1xPBS (Lonza) with 1/1000 diluted cisplatin (Fluidigm). Cells were then incubated for 1h at RT (or left at 4°C overnight) in the Iridium-intercalator solution in fixation and permeabilization buffer (BD Cytofix/Cytoperm™, BD Biosciences). Then cells were washed with 1x permeabilization buffer (BD Perm/Wash™, BD Biosciences) and stained for 30min at RT with intracellular antibody cocktail diluted in 1x permeabilization buffer (**Table S17**). Cells were fixed with BD Cytofix/Cytoperm™ and left overnight until analysis, or immediately used for mass cytometry. Immediately before starting the analysis, cells were washed in CyFACS buffer, then PBS and finally with MiliQ water. Prior to data acquisition, cells were resuspended in MilliQ water containing 1/10 diluted normalization beads (EQ Four Element Calibration Beads, Fludigm) to the concentration of less than 8×10^5^ cells/ml to achieve an acquisition rate of 400 events/s on the CyTOF Helios mass cytometer (Fluidigm). In each sample 1-1.5 million cells were acquired. After acquisition, data were normalized with the reference EQ passport P13H2302 (70) and further data analysis was performed using FlowJo v10.

### Statistical Analysis

All the statistical parameters (sample size, statistical tests, and statistical significance) are reported in the Figures and Figure Legends. Significance of differences in frequencies of the immune cell subsets between high and low responders in the datasets was calculated using the Significance analysis of microarrays (SAM) (71) at false discover rate (FDR) < 1%. Mass cytometry data between two groups after peptide stimulation were analyzed using the one-way ANOVA Kruskal-Wallis test followed by Dunn’s multiple comparison test, while paired samples within groups were compared with two-tailed Wilcoxon matched-pairs signed rank test. Additionally, pairwise t-test with the Benjamini-Hochberg (B-H) correction for multiple testing adjustment with 0.95 confidence level was used to evaluate changes in the cell frequencies after vaccination within groups. Pearson’s correlation coefficient was used to evaluate the correlations between features from the top-performing models. The Corrplot package in R was used to calculate correlation coefficients, statistics and for visualization of the correlation matrix (72). P-values were adjusted for multiple comparisons by using the Benjamini-Hochberg correction (73). Statistical analyses were performed with GraphPad PRISM 7.04 (Graph Pad Software) or in R, and p values above 0.05 were considered not significant.

### Code and data availability

The source code of the *mulset* algorithm is available from https://github.com/LogIN-/mulset. The *mulset* is available as an R package in CRAN, a repository of open-source software. Pseudo-code for SIMON is available as **Data file S1**. All data used in SIMON analysis are available from the Stanford Data Miner (www.datamt.net). Mass cytometry fcs files related to Figure 4 (https://zenodo.org/record/1328286) are available on a research data repository Zenodo maintained by OpenAIRE and CERN (www.zenodo.org).

## Supporting information

Pseudo-code for SIMON

## Supplementary Materials

Fig. S1. Distribution of high and low responders included in the initial dataset based on gender, CMV status and study year

Fig. S2. Assays performed across different clinical studies and study years.

Fig. S3. Staining profiles and gating scheme of immune cell subsets analyzed using mass cytometry.

Fig. S4. Visualization of the initial dataset in the context of missing values.

Fig. S5. Performance evaluation of models build on datasets 13 and 36 after applying restriction filters.

Fig. S6. Heatmap of the correlation coefficients calculated between features from the dataset 13.

Fig. S7. Importance of features determined by Boruta.

Fig. S8. Heatmap of the correlation coefficients calculated between features from the dataset 36.

Table S1. List of 102 analyzed immune cell subsets showing gating strategy.

Table S2. Immune cell subsets and phosphorylation of proteins identified using phosphorylated cytometry.

Table S3. Cytokines, chemokines and growth factors analyzed by Luminex. Table S4. Sparsity calculated by column.

Table S5. 34 datasets generated using intersections.

Table S6. List of machine learning algorithms implemented in SIMON.

Table S7. List of all models built and their minimal and maximal AUROC values.

Table S8. List of all models with minimal and maximal AUROC values after applying performance restriction filters.

Table S9. List of models with maximal train and test AUROC.

Table S10. All models built on dataset 36 after restriction filters applied.

Table S11. All models built on dataset 13 after restriction filters applied.

Table S12. Characteristics of individuals with high and low response to influenza vaccination selected in the dataset 13.

Table S13. List of features and their variable importance score in dataset 13.

Table S14. Characteristics of individuals with high and low response to influenza vaccination used for experimental validations.

Table S15. Characteristics of individuals with high and low response to influenza vaccination selected in the dataset 36.

Table S16. List of features and their variable importance score in dataset 36. Table S17. Antibody panel for ICS mass cytometry.

Table S18. List of peptides in the influenza peptide pool. Data file S1. Pseudocode for SIMON.

## Acknowledgments

We are grateful to all individuals that participated in the research studies. Special acknowledgment to Dr. Purvesh Khatri for critical reading the manuscript. We appreciate helpful discussions and support from all members of the Davis and Y. Chien labs, specifically Elsa Sola, Allison Nau, Lisa Wagar and Asbjorn Christophersen for help with mass cytometry and input from Paula Romer. We also thank all staff members from the HIMC (Michael D. Leipold) for data analysis, management and helpful discussions and HIMC Biobank (Rohit Gupta and Janine Bodea Sung) for sample processing and storage, Stanford-LPCH Vaccine Program (Alison Holzer) for management of clinical studies and Stanford FACS facility for all the support.

## Funding

This work was supported by NIH grants (U19 AI090019, U19 AI057229) and the Howard Hughes Medical Institute to M.M.D, and by the EU’s Horizon 2020 research and innovation program under the Marie Sklodowska-Curie grant (FluPRINT, Project No 796636) to A.T.

## Author Contributions

A.T. designed and performed the experiments, processed and analyzed the data and wrote the manuscript. I.T. designed the database, programmed the SIMON, analyzed the data and revised the manuscript. Y.R.H. and H.T.M. run all the experiments at the HIMC, analyzed the data and revised the manuscript. C.L.D. was responsible for regulatory approvals, protocol design, study conduct, and clinical data management. M.M.D. supervised the study and edited the manuscript.

## Conflict of interests

The authors declare that they have no conflict of interests.

## Data and materials availability

The source code of the mulset algorithm is available from https://github.com/LogIN-/mulset. The *mulset* is available as an R package in CRAN, a repository of open-source software. Pseudo-code for SIMON is available as **Data file S1**. All data used in SIMON analysis are available from the Stanford Data Miner (www.datamt.net). Mass cytometry fcs files related to Figure 4(https://zenodo.org/record/1328286) are available on a research data repository Zenodo maintained by OpenAIRE and CERN (www.zenodo.org).

