## Supplementary figures S1-S8 for "SIMON, an automated machine learning system reveals immune signatures of influenza vaccine responses"

### Supplementary Materials:

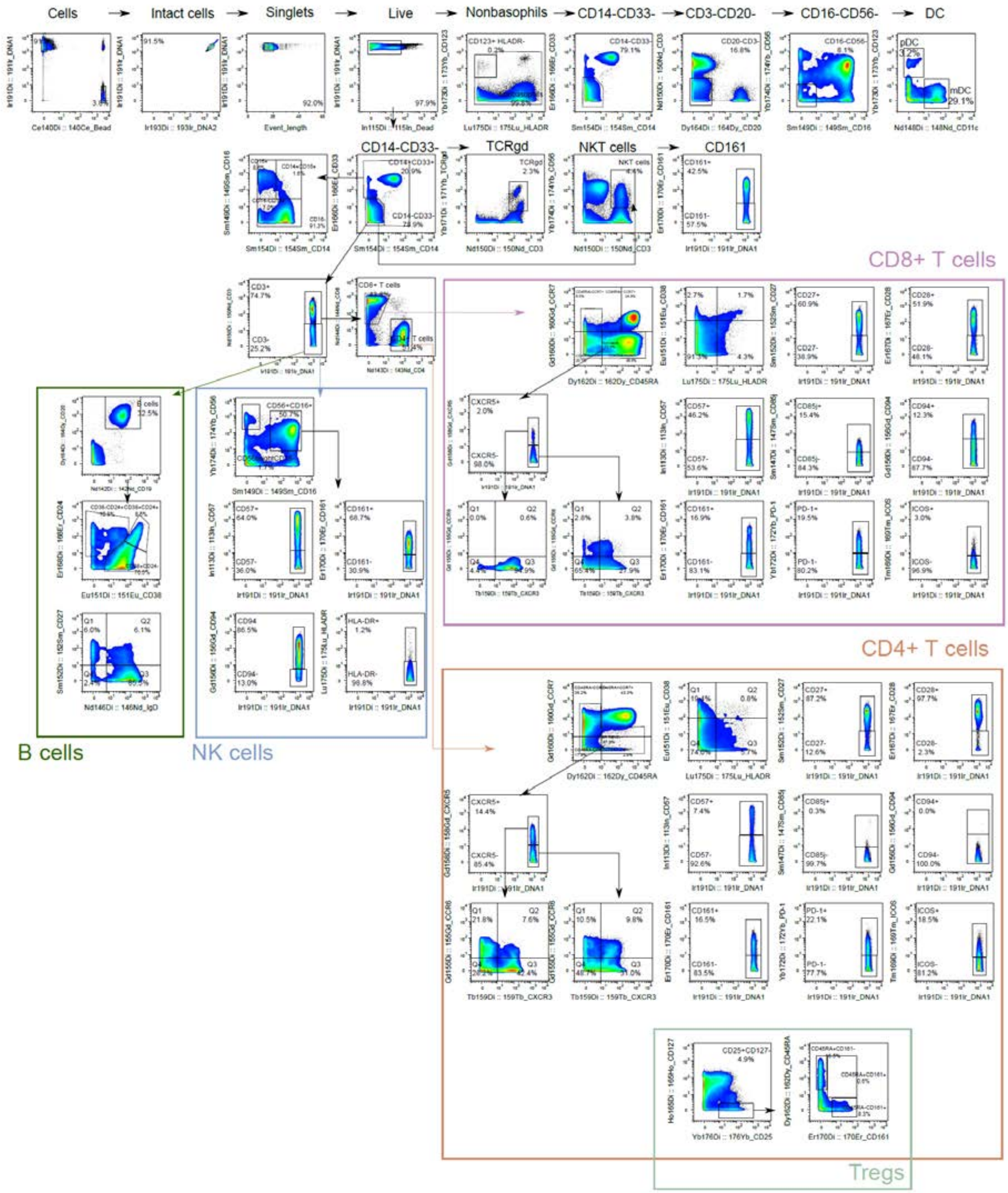

**Figure S1. Staining profiles and gating scheme of immune cell subsets analyzed using mass cytometry.** Representative gating strategy for phenotype analysis of different blood-derived immune cell subsets analyzed using mass cytometry in the sample from one donor acquired before vaccination. In total PBMC from healthy 187 donors were analyzed using same gating scheme. Text above plots indicates parent population, while arrows show gating strategy defining major immune cell subsets (CD4+ T cells, CD8+ T cells, B cells, NK cells, Tregs, NKT cells, etc.).

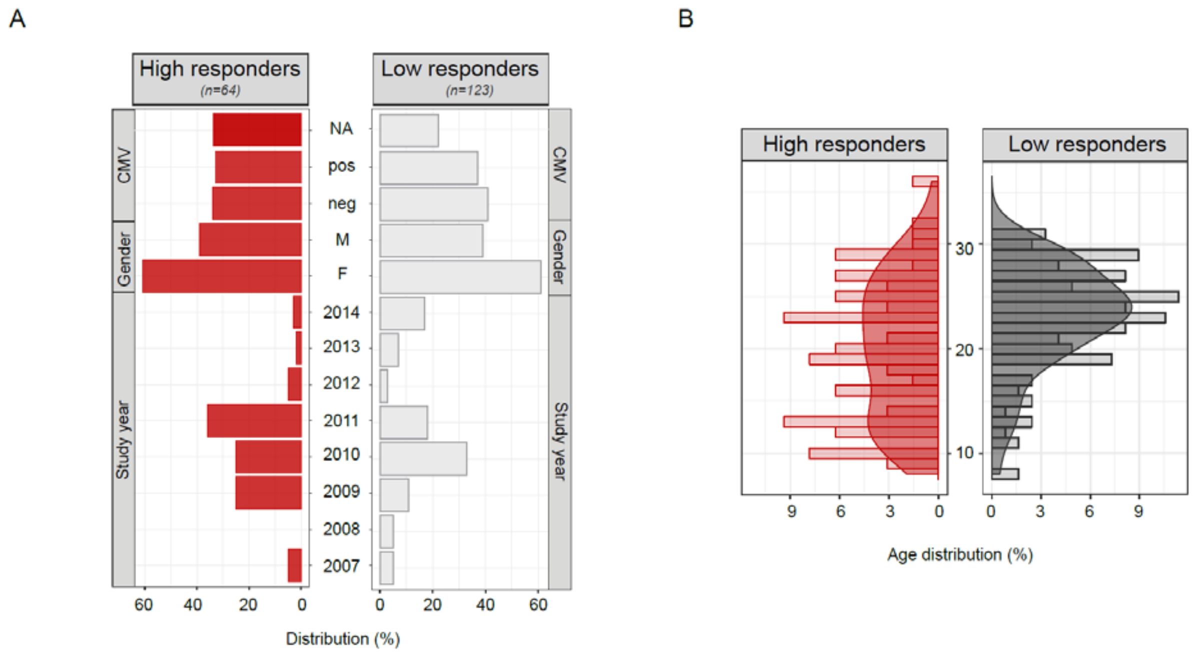


**Figure S2. Distribution of high and low responders included in the initial dataset.** Distribution of individuals in groups of high (red, n=64) and low (grey, n=123) responders regarding the (**A**) CMV status, gender and study year. (**B**) Age distribution between high and low responders. Age is indicated in years.

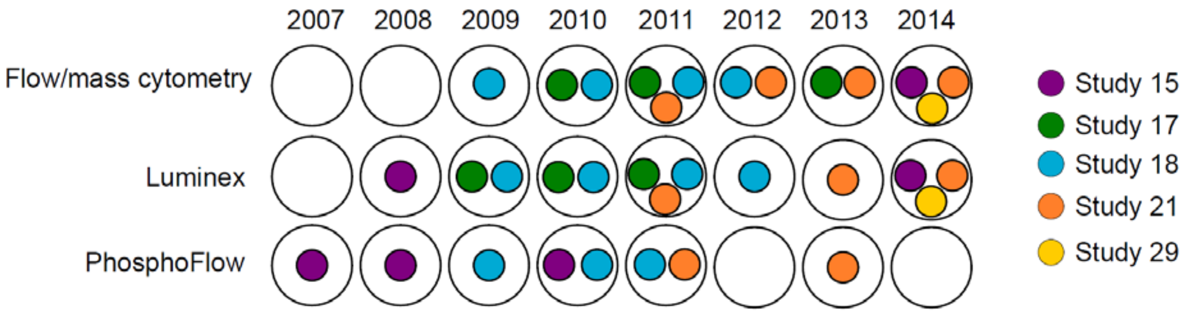


**Figure S3. Assays performed across different clinical studies and study years.** Data from 5 different clinical studies (Study 15, 17, 18, 21 and 29) were included in the analysis. Flow cytometry was performed only in year 2009, in other years phenotype of immune cells was determined by mass cytometry. Luminex (either 51/63-plex) was performed from 2008 to 2014. Finally, signaling capacity of immune cells was analyzed by phosphorylation cytometry (PhosphoFlow) on mass cytometer in 2013 and flow cytometer in all other years.

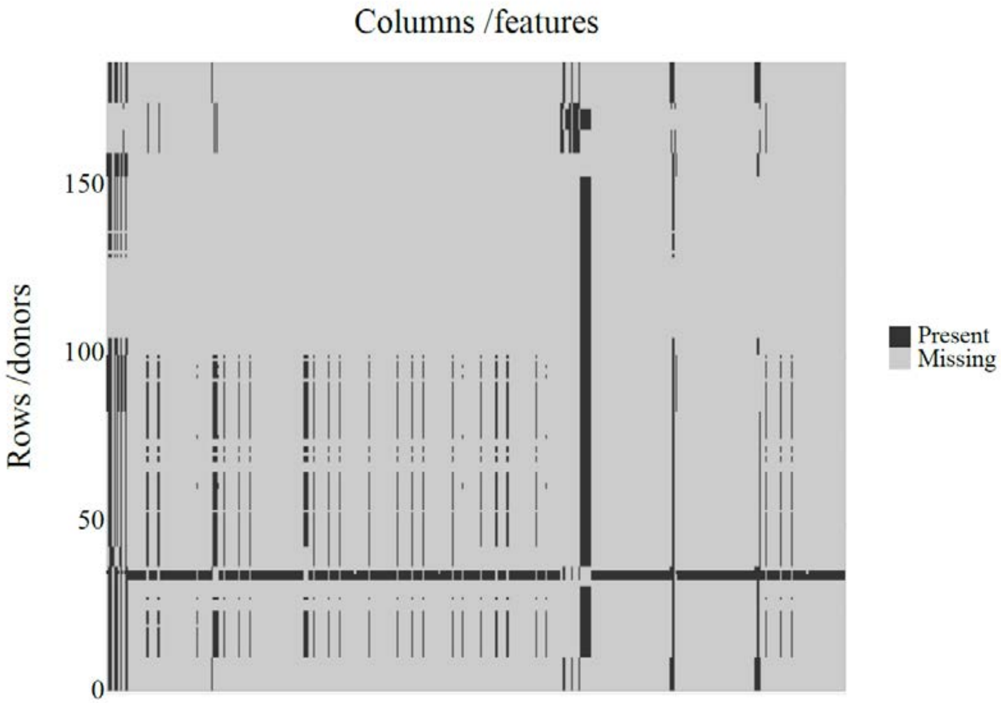


**Figure S4. Visualization of the initial dataset in the context of missing values.** Heatmap showing distribution of data in the initial dataset. Each row represents a unique donor, while each column is one feature. Missing values are shown in grey, while present values are shown in black.

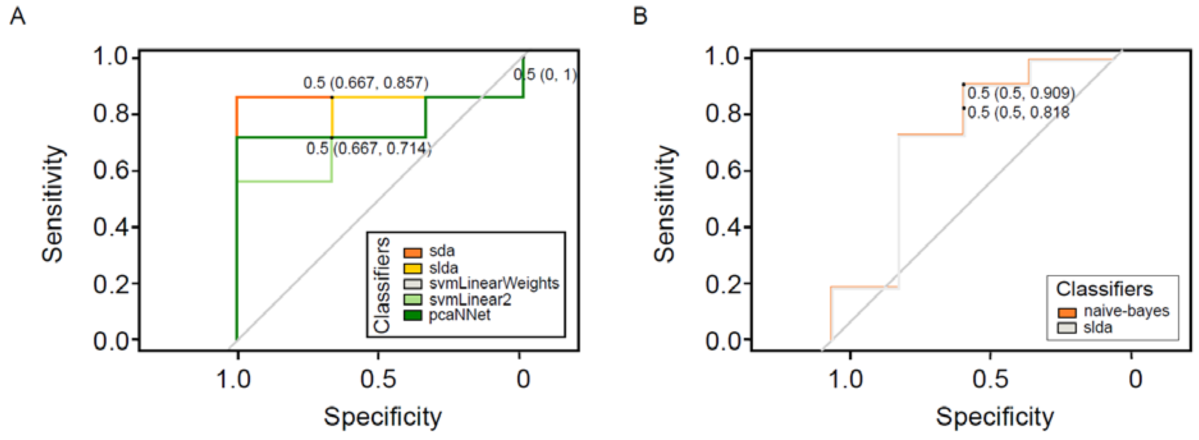


**Fig. S5. Performance evaluation of models build on datasets 13 and 36 after applying restriction filters.** ROC curves shown for all the models build on **(A)** dataset 36 and **(B)** dataset 13. Each model (classifier) is denoted in the color indicated in the graph legend.

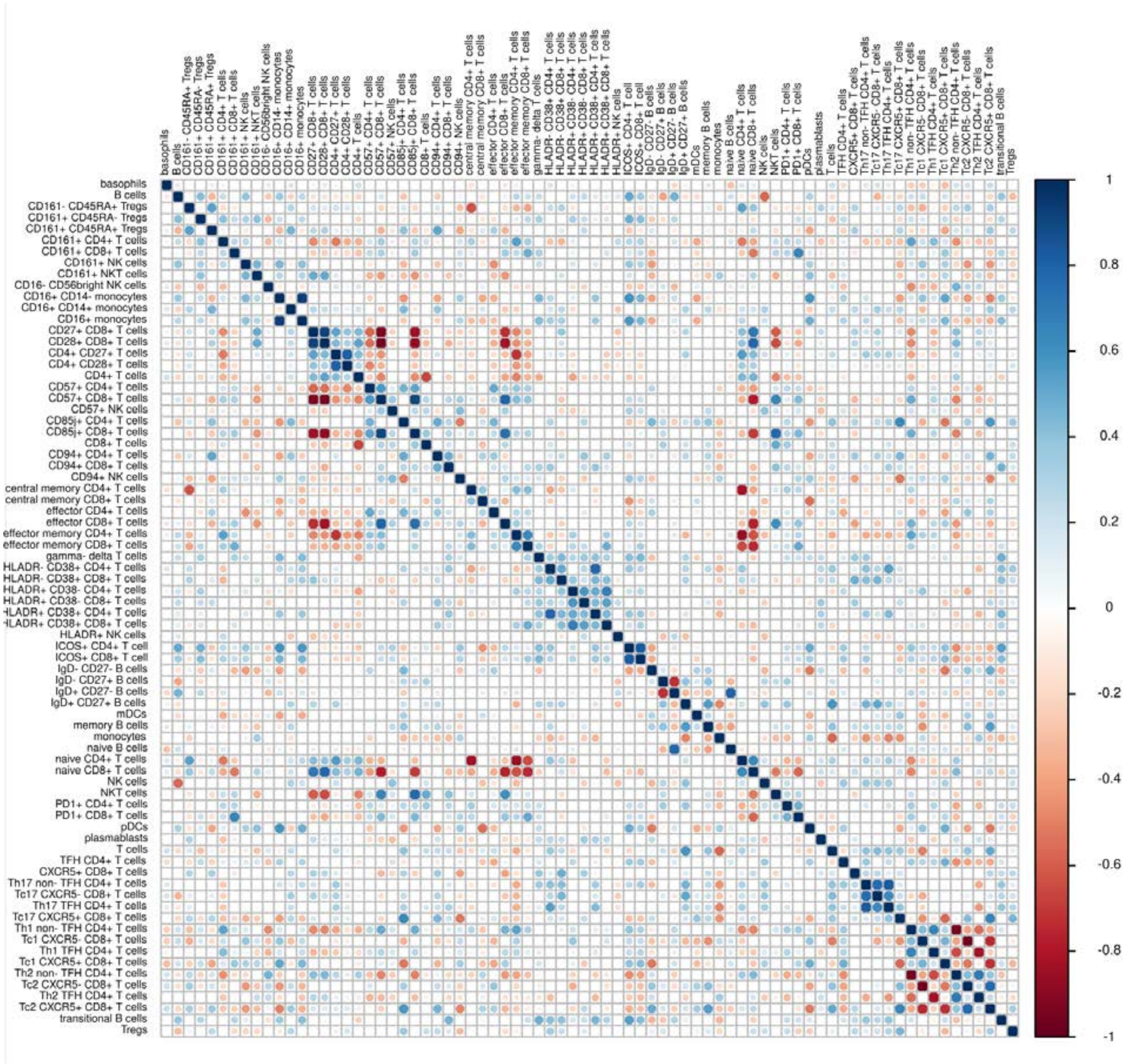


**Fig. S6. Heatmap of the correlation coefficients calculated between features from the dataset 13.** Heatmap shows the significant correlation coefficients between all the features from dataset 13 calculated using Pearson correlation ($p<0.05$). Not significant values are shown as blank. Color of each circle follows the legend on the right side of the heatmap and red indicates values with negative correlation, while blue values with positive correlation.

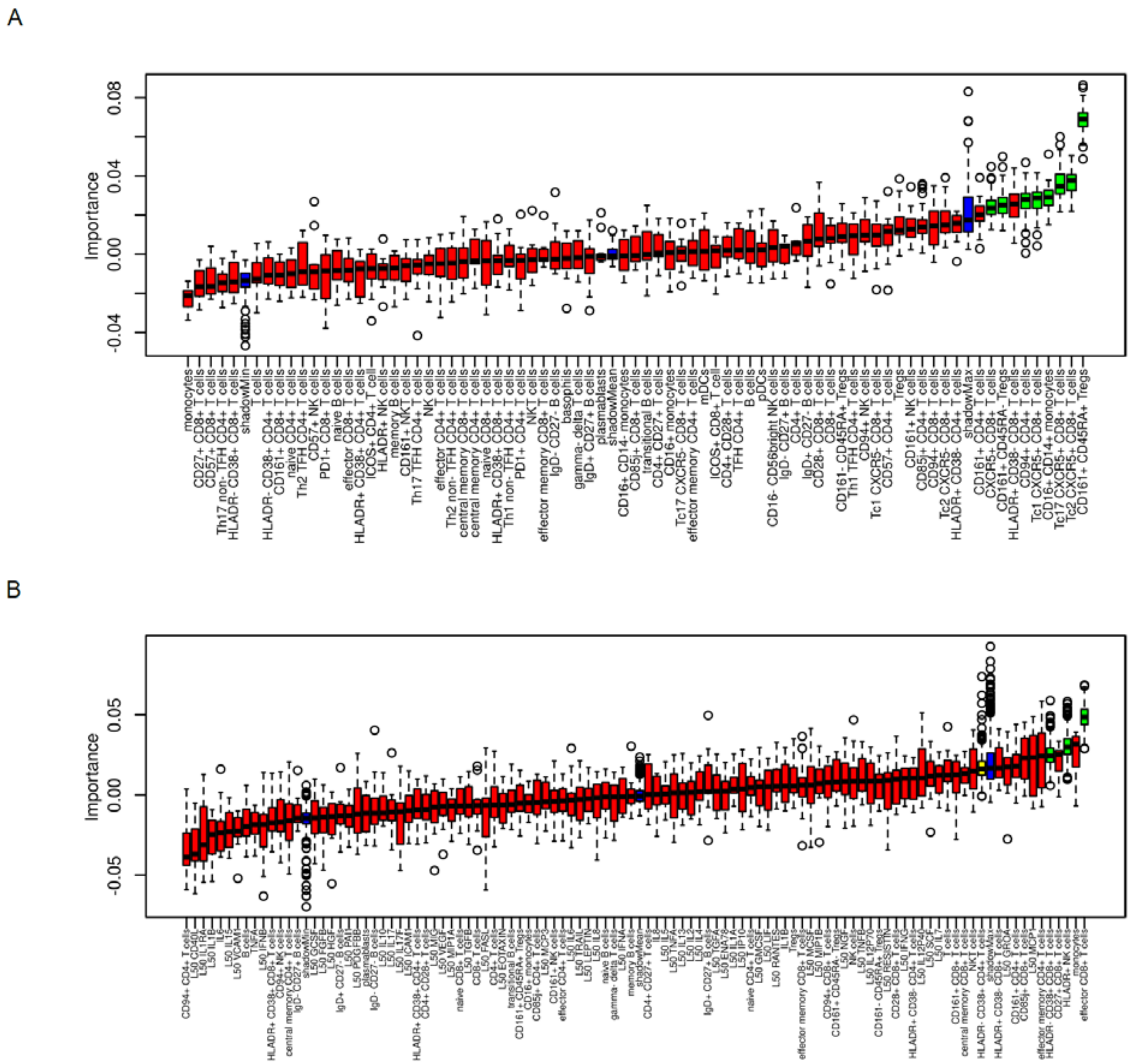


**Fig. S7. Importance of features determined by Boruta.** Boruta result plots for **(A)** dataset 13 and **(B)** dataset 36. Red boxplots represent importance score of rejected features, while green boxplots show minimal, average and maximum importance score for confirmed features. Blue boxplots show importance score of a shadow feature. Yellow boxplots are tentative features.

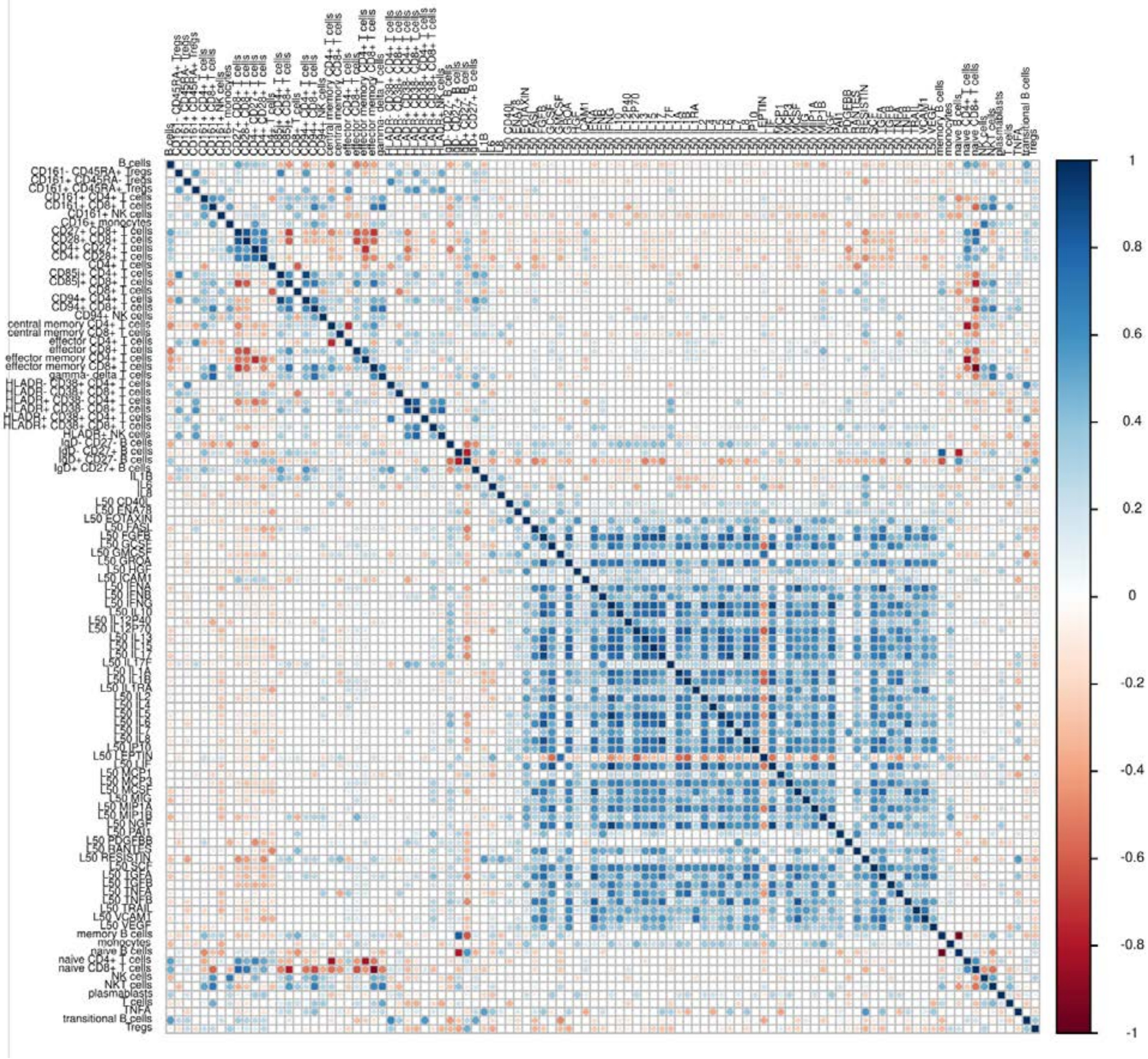


**Fig. S8. Heatmap of the correlation coefficients calculated between features from the dataset 36.** Heatmap shows the significant correlation coefficients between all the features from dataset 36 calculated using Pearson correlation ($p<0.05$). Not significant values are shown as blank. Color of each circle follows the legend on the right side of the heatmap and red indicates values with negative correlation, while blue values with positive correlation.

**Tables S1-S19 provided in single Excel file**

**Data files S1-S2**

### Online Methods References

**List of references of R packages used for Supplementary Table 6:**

12. Algorithm: bagFDA Package: earth - Stephen Milborrow. Derived from mda:mars by Trevor Hastie and Rob Tibshirani. Uses Alan Miller's Fortran utilities with Thomas Lumley's

leaps wrapper. (2017). earth: Multivariate Adaptive Regression Splines. R package version 4.5.0. https://CRAN.R-project.org/package=earth

44. Algorithm: gpls Package: gpls - Beiying Ding (2017). gpls: Classification using generalized partial least squares. R package version 1.48.0.

54. Algorithm: logicBag Package: logicFS - Holger Schwender (2013). logicFS: Identification of SNP Interactions. R package version 1.46.0.

82. Algorithm: plsRglm Package: plsRglm - Frederic Bertrand, Nicolas Meyer and Myriam Maumy-Bertrand (2014). Partial Least Squares Regression for Generalized Linear Models, R package version 1.1.1

101. Algorithm: RRFglobal Package: RRF - H. Deng(2013). Guided Random Forest in the RRF Package. arXiv:1306.0237.

124. Algorithm: vglmAdjCat Package: VGAM - Thomas W. Yee (2015). Vector Generalized Linear and Additive Models: With an Implementation in R. New York, USA: Springer.

125. Algorithm: vglmContRatio Package: VGAM - Thomas W. Yee (2015). Vector Generalized Linear and Additive Models: With an Implementation in R. New York, USA: Springer.

126. Algorithm: vglmCumulative Package: VGAM - Thomas W. Yee (2015). Vector Generalized Linear and Additive Models: With an Implementation in R. New York, USA: Springer.

127. Algorithm: widekernelpls Package: pls - Bjørn-Helge Mevik, Ron Wehrens and Kristian Hovde Liland (2016). pls: Partial Least Squares and Principal Component Regression. R package version 2.6-0. https://CRAN.R-project.org/package=pls

128. Algorithm: wsrf Package: wsrf - He Zhao, Graham J. Williams, Joshua Zhexue Huang (2017). wsrf: An R Package for Classification with Scalable Weighted Subspace Random Forests. Journal of Statistical Software, 77(3), 1-30. doi:10.18637/jss.v077.i03

**Other R packages used**

1. Hadley Wickham (NA). feather: R Bindings to the Feather 'API'. R package version 0.3.1. https://github.com/wesm/feather
2. Package: RMySQL - Jeroen Ooms, David James, Saikat DebRoy, Hadley Wickham and Jeffrey Horner (2017). RMySQL: Database Interface and 'MySQL' Driver for R. R package version 0.10.11. https://CRAN.R-project.org/package=RMySQL
3. Package: DBI - R Special Interest Group on Databases (R-SIG-DB), Hadley Wickham and Kirill Müller (2017). DBI: R Database Interface. R package version 0.7. https://CRAN.R-project.org/package=DBI
4. Package: dplyr - Hadley Wickham, Romain Francois, Lionel Henry and Kirill Müller (2017). dplyr: A Grammar of Data Manipulation. R package version 0.7.1. https://CRAN.R-project.org/package=dplyr
5. Package: plyr - Hadley Wickham (2011). The Split-Apply-Combine Strategy for Data Analysis. Journal of Statistical Software, 40(1), 1-29. URL http://www.jstatsoft.org/v40/i01/
6. Package: reshape2 - Hadley Wickham (2007). Reshaping Data with the reshape Package. Journal of Statistical Software, 21(12), 1-20. URL http://www.jstatsoft.org/v21/i12/

7. Package: doMC - Revolution Analytics and Steve Weston (2015). doMC: Foreach Parallel Adaptor for 'parallel'. R package version 1.3.4. https://CRAN.R-project.org/package=doMC

8. Package: jsonlite - Jeroen Ooms (2014). The jsonlite Package: A Practical and Consistent Mapping Between JSON Data and R Objects. arXiv:1403.2805 [stat.CO] URL https://arxiv.org/abs/1403.2805
