## Supplementary material for "SIMON, an automated machine learning system reveals immune signatures of influenza vaccine responses": Pseudo-code for SIMON

```
Gather user input: List of csv files (i.e. in this work 177 csv files collected from Stanford HIMC)

% Step 1: data import
for {each csv file to be analyzed} do:
    Read data matrix from one csv file, remove irrelevant csv columns, standardize and sanitize row values (i.e. standardize identifiers/values inconsistency)
    Impute data (i.e. calculate vaccine response variable such as GMT HAI titer)
end for;
Import newly created data into the MySQL database

% Step 2: select data
Selection of data based on the research question (i.e. inclusion criteria for donors)

% Step 3: generate re-sampled intersection datasets suitable for analysis
for {each subject in data} do:
    Calculate intersection between subject and all other subjects using mulset algorithm
    Skip sets that have less than 5 features in common
end for;
Save all shared intersections to corresponding datasets

% Step 4: automated machine learning
avaliableModels – install machine learning R packages necessary for building models (here, we selected 128 classification algorithms)
for {dataset in sets} do:
    Create balanced partitioning of the data
    data: 75% training, 25% test
    Skip dataset if test set has less than 10 subjects
    for {model in avaliableModels} do:
        Perform model training and get all model performance variables
        Using test data make predictions on the trained model, retrieve ROC from confusion matrix
        Using trained model calculate variable importance score
        Save all data metrics to corresponding fields in the database
    end for;
end for;
```
